# A resurrection experiment finds evidence of both reduced genetic diversity and potential adaptive evolution in the agricultural weed *Ipomoea purpurea*

**DOI:** 10.1101/024950

**Authors:** Adam Kuester, Ariana Wilson, Shu-mei Chang, Regina S. Baucom

**Affiliations:** 2059 Kraus Natural Science Building Department of Ecology and Evolutionary Biology 830 North University University of Michigan Ann Arbor, Michigan, 48109; 2502 Plant Sciences Building Plant Biology Department University of Georgia 120 Carlton StreetAthens, GA 30602

**Keywords:** agriculture, allelic diversity, selection, genetic drift, temporal evolution

## Abstract

Despite the negative economic and ecological impact of weeds, relatively little is known about the evolutionary mechanisms that influence their persistence in agricultural fields. Here, we use a resurrection ecology approach and compare the genetic and phenotypic divergence of temporally sampled seed progenies of *Ipomoea purpurea*, an agricultural weed that is resistant to glyphosate, the most widely used herbicide in current-day agriculture. We found striking reductions in allelic diversity between cohorts sampled nine years apart (2003 vs 2012), suggesting that populations of this species sampled from agricultural fields have experienced genetic bottleneck events that have led to lower neutral genetic diversity. Heterozygosity excess tests indicate that this bottleneck may have occurred prior to 2003. Further, a greenhouse assay of individuals sampled from the field as seed found that populations of this species, on average, exhibited modest increases in herbicide resistance over time. Our results show that populations of this noxious weed, capable of adapting to strong selection imparted by herbicide application, may lose genetic variation as a result of this or other environmental factors. We likely uncovered only modest increases in resistance between sampling cohorts due to a strong and previously identified fitness cost of resistance in this species, along with the potential that non-resistant migrants germinate from the seed bank.

## Introduction

The influence of human mediated selection is perhaps nowhere more prevalent than in the agricultural system. Agricultural weeds, in particular, provide excellent case studies of adaptation to human-mediated selection (Baker 1974). They are exposed to fertilizers, herbicides, irrigation, as well as variable cropping techniques, and these manipulations can impose frequent, strong, and highly predictable disturbance regimes (Barrett 1988). Examples of rapid adaptation to these scenarios are present in the literature from early cases of crop mimicry (Baker 1974; Barrett 1983; Barrett 1988) to the many recent examples of the evolution of herbicide resistance (Heap 2016). Although weeds are in many cases models for understanding rapid evolution and persistence in stressful environments, we currently have a limited understanding of the broad genetic changes that may influence weed populations in agricultural landscapes (Vigueira *et al*. 2013; Waselkov & Olsen 2014). These lapses in our knowledge are striking because the population dynamics of agricultural weeds are directly relevant to the global food supply. Agricultural weed infestations reduce world-wide crop yield by as much as 10% (Oerke 2005) and it has been estimated that crop losses caused by weeds cost the US agricultural economy ~33B USD per year (Pimentel *et al*. 2005). Clarifying the evolutionary forces that impact agricultural weeds can provide information on the process of rapid evolution more broadly as well as insight on how weeds survive and persist in agricultural regimes.

Although agricultural weeds co-exist and compete with crops, they evolve though unintentional human mediated selection rather than through direct artificial selection (Stewart & Warwick 2005) and represent a novel evolutionary state that is “neither wild nor domesticated” (Vigueira *et al*. 2013). Weeds are subject to the same forces influencing evolution in nature— notably, genetic drift, selection, and gene flow (Jasieniuk *et al*. 1996)—but they often experience a selection intensity that is much higher than what is usually found in other natural systems. For example, the predominant form of weed control in current farming is through the use of herbicides, which are designed to remove 90% of the weed population (Jasieniuk *et al*. 1996; Délye *et al*. 2013). Individuals that survive this high intensity of selection due to either chance or genetic predisposition are founders for the next generation. Since the point of weedy plant control regimes—whether through the use of herbicide or another control technique—is to remove of a large portion of the population, populations that re-colonize are hypothesized to show a pattern of constriction and genetic bottleneck (Jasieniuk *et al*. 1996; Vigueira *et al*. 2013). As a result, weeds could lose rare alleles important to future adaptation (Nei *et al*. 1975).

In support of this idea, population genetic surveys have found that weeds tend to exhibit less genetic variation than other groups of plants (Hamrick *et al*. 1979), and there is some evidence that weed populations from cultivated land exhibit decreased neutral genetic diversity compared to wild populations (Kane & Rieseberg 2008). The majority of the work to date, however, has compared populations across space, *i.e.,* from cultivated and non-cultivated areas (Muller *et al*. 2010), or “wild” versus “weedy” populations (Kane & Rieseberg 2008). In contrast, a novel approach that can provide direct evidence for evolutionary change through time is by the use of “resurrection ecology” in which ancestor and descendant strains of species are compared. In this type of experiment, seeds or propagules sampled from an earlier time point are germinated after remaining dormant for a number of years and compared to descendant populations sampled from the same location (Franks *et al*. 2007; Orsini *et al*. 2013). Although resurrection ecology experiments have been used to address key questions on evolutionary constraints in microbial systems (Lenski & Travisano 1994; Lenski 1998), such experiments in eukaryotes have thus far used either a limited number of accessions (Baucom & Mauricio 2010) or a limited number of distinct populations (Franks *et al*. 2007; Thomann *et al*. 2015).

Here we perform a resurrection ecology experiment to examine the population genetics and potential adaptability of *Ipomoea purpurea,* the common morning glory. This species is native to the central highlands of Mexico (Clegg & Durbin 2000; Defelice 2001) and is an introduced invader of agricultural and disturbed areas in the United States (Defelice 2001). Although the precise history of introduction into the US is unknown (Fang *et al*. 2013), lineages in the US exhibit low diversity relative to Mexican accessions, suggesting a severe bottleneck occurred following introduction (Fang *et al*. 2013). *I. purpurea* is considered one of the worst weeds of southeastern US agriculture (Baucom *et al*. 2011), in part because it exhibits low-level resistance to glyphosate, the most utilized herbicide in agriculture world-wide (Powles 2008). Recently, we have identified a mosaic of glyphosate resistance across the US, with some populations of *I. purpurea* exhibiting high resistance (a high proportion of the population that survives glyphosate) and others showing high susceptibility post-herbicide application (Kuester *et al*. 2015). Previous work has also found that an additive genetic basis underlies glyphosate resistance in this species (Baucom & Mauricio 2008) and that resistance segregates in genetic lines developed from a single population (Debban *et al*. 2015). Although *I. purpurea* populations are found primarily within agricultural fields that are treated with glyphosate and other herbicides, the impact of such strong selection, and any associated environmental changes on the population genetics of this species over time remains largely unknown. Because seeds remain viable for many years in lab conditions, we are able to examine both neutral and adaptive genetic variation of populations sampled from the same agricultural fields at two different time points—once in 2003 and again in 2012 (see Figure 1 for population locations). We first determine if the neutral genetic differentiation and diversity of *I. purpurea* populations have changed between sampling years. We pair this with greenhouse experiments to examine the potential that these populations, sampled from the same fields that were used for either soy or corn farms between 2003 and 2012 (Table S1) exhibit increased resistance over time. We find evidence of both genetic bottleneck and slight increase in the level of resistance, indicating that a noxious weed potentially adapts to the extreme selection imposed by herbicide applications even as genetic diversity decreases. We further find some indication that highly resistant populations exhibit lower genetic diversity than less resistant populations, suggesting that herbicide application is responsible for the reduction in neutral genetic diversity. This is the first examination, to our knowledge, of a resurrection ecology experiment that simultaneously identifies both loss of genetic diversity of an agricultural weed over time as well as potential evidence for adaptive evolution.

**Figure 1.**
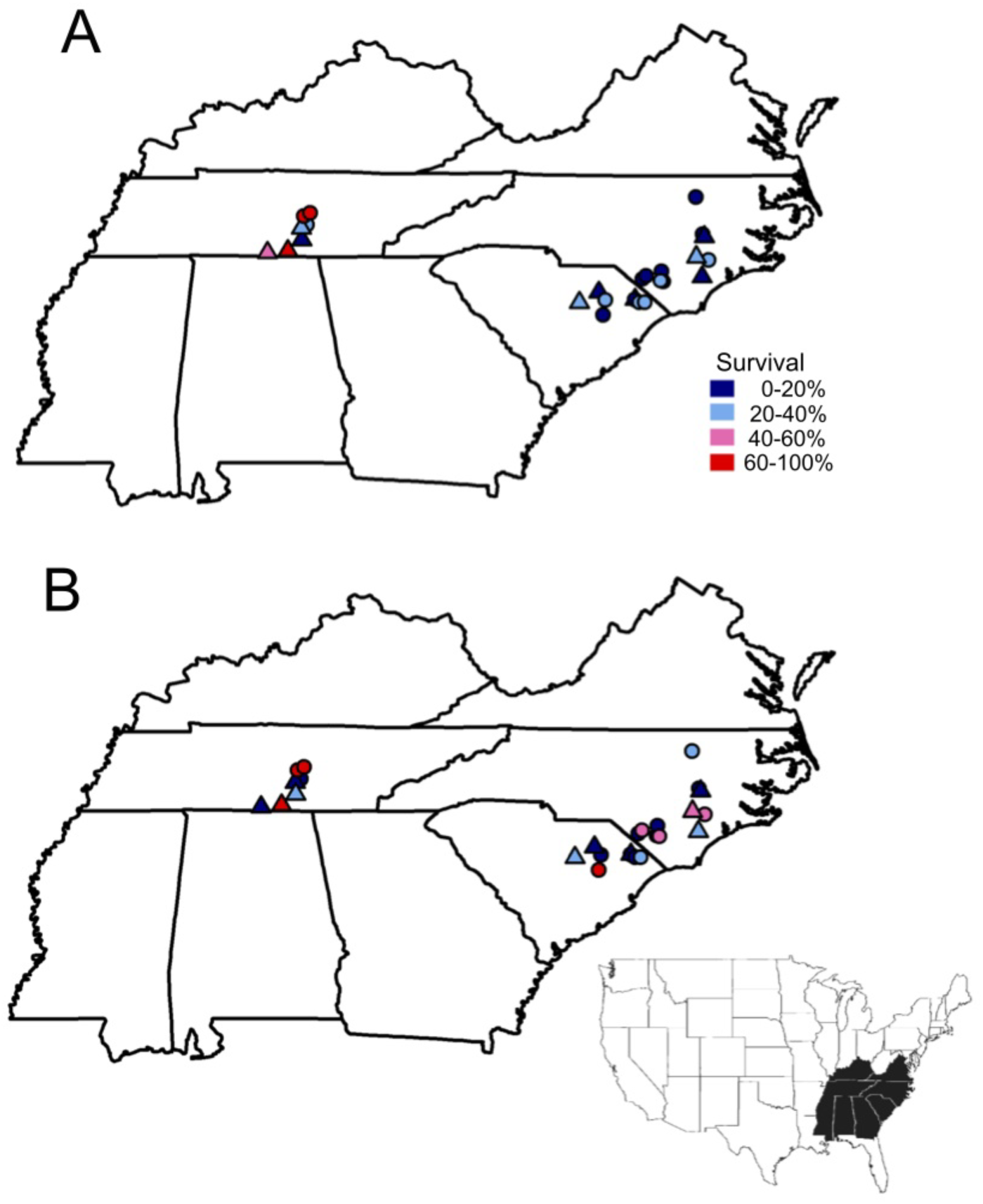
Map of populations sampled from A) 2003 and B) 2012 within the US. Populations that were genotyped in both 2003 and 2012 are indicated by a triangle (see Table S1 for sites used for resistance measurements). The percent survival following 3.4 kg ai/ha of RoundUp is indicated in color. Sites were sampled at least 5 km apart.

## Materials and Methods

**Population sampling.** Locations and sampling strategies for 44 *I. purpurea* populations were previously described in Kuester *et al*. (2015). Twenty-six of these populations were sampled in 2003 and resampled in 2012 (see Figure 1 and Table S1). In 2003, we collected replicate seeds from between 6-30 maternal individuals at least 1 m apart from one another along a linear transect. We located the same populations in the fall of 2012 using GPS coordinates, which are accurate to within a few meters. Agricultural fields are highly disturbed by tilling and harvesting each year, and morning glories are predominantly found in areas that have recently experienced soil disturbance *via* tilling; as a result, this system is not amenable to the maintenance of longterm transects. We are thus making the assumption that adult plants present within the same agricultural field, and located within the nearest distance to the GPS coordinates in 2012 are the descendants of the 2003 cohort. Preliminary data from >5,000 SNPs generated by genotype-by-sequencing has identified a high number of independent genetic clusters in population structure analyses and a low proportion of recent immigrants into populations (Alvarado-Serrano *et al*. unpublished data) indicating that our assumptions herein are largely realistic. We estimated population size in the 2012 sampling year by counting the numbers of individuals down a linear transect.

Of the 26 populations that were sampled both years, we randomly chose 10 to examine potential changes in genetic diversity between 2003 and 2012. One seed from an average of 18 maternal lines per population per sampling year (355 individuals total) were germinated and cotyledons were used for DNA isolation using a CTAB method modified from Stokes et al. 2009 (see Kuester *et* al.2015). The numbers of maternal lines sampled per population were approximately equal between the sampling years and exact numbers are presented in Table S2.

To assay herbicide resistance among populations and between sampling years, we planted two replicate greenhouse experiments of all 26 populations at the University of Georgia Plant Biology Greenhouses (Athens, GA). One seed from 10 maternal lines per population per sampling year were scarified and planted in pine bark soil in SC10 super conetainers (Stuewe and Sons, Tangent, OR) in six experimental treatments, described below. This design was replicated in its entirety in another greenhouse for a total of 20 seeds per population within each treatment and thus an overall total of 5381 experimental individuals. Plants were randomly assigned to racks that were then randomly assigned to flow trays (4 racks per flow tray). Conetainers were watered daily and flow trays were filled with water to prevent desiccation. Germination was slightly higher in 2003 compared to 2012 samples (87% and 84% in 2003 and 2012, respectively, χ^21^ = 12.27, P < 0.001) and ranged from 50-98% across populations.

Plants were sprayed with RoundUp PowerMax (Monsanto, St Louis, MO) 22 days after planting at rates around the recommended field rate (1.54 kg ai/ha) of 0, 0.21, 0.42, 0.84, 1.70 and 3.40 kg a.i./ha (the 0 kg a.i./ha control treatment was sprayed with water) using a hand-held, CO_2_ pressurized sprayer (R & D Sprayers, Opelousas, LA) that delivered 187 L ha^−1^ at 206 kPa, 1.5 meters above the plants. Three weeks after glyphosate application we scored survival of each plant. Plants were harvested, dried at 72°C for 48 hours and measured for total above ground biomass. Biomass values were adjusted to the non-sprayed controls by dividing each individual by the average biomass of its population grown in the non-spray control treatment following standard protocols (Tehranchian *et al*. 2015).

**SSR genotyping and scoring errors.** Details on multiplexing SSR markers and scoring procedures can be found in Kuester *et al*. (2015). Briefly, 15 polymorphic microsatellite loci were used to examine genetic diversity across populations and sampling years, and all individuals were scored by hand. To check accuracy of multi-locus genotypes we re-scored loci from 200 randomly chosen individuals and found very few scoring errors. We did not find any large allele drop-outs or errors due to stutter in any of the locus by population by year combinations. We also examined the influence of null alleles on genetic diversity and found little evidence that potential null alleles altered our estimates or the main conclusions. Details of these analyses are presented in the Supporting Information section.

**Temporal genetic differentiation and diversity**. We examined the potential that seeds sampled across collection years were genetically differentiated from one another in two ways. First, we estimated genetic differentiation between years (FRT) using hierarchical AMOVA in GenAlEx v. 6.5 (Peakall & Smouse 2012). We also performed individual assignment (Paetkau *et al*. 1995; Cornuet *et al*. 1999) of individuals to sampling year using GeneClass2 (Piry *et al*. 2004). For individual assignment, the inability to assign individuals to a specific sampling year would indicate that individuals sampled in 2012 had not diverged in allelic composition compared to the individuals sampled in 2003. We used the Bayesian method described by Baudouin and Lebrun (Baudouin & Lebrun 2000) as a criterion for computation, and individual assignment was performed using the leave-one-out procedure (Paetkau *et al*. 2004), where the genotype to be assigned was not included in the population from which it was sampled. We report the - log likelihood of being assigned in each sampled year, by plotting the - log likelihood value of individual assignment to 2003 sample year against the - log likelihood of being assigned to the 2012 sampling year. Lack of temporal change across sampling years would be indicated by overlap of individuals sampled from each year. We calculated expected and observed heterozygosity (He and Ho), the number of alleles (Na) and the number of effective alleles (Ne) using GenalEx v 6.5 (Peakall & Smouse 2012) and and allelic richness (AR) using FSTAT v. 2.9.3.2 (Goudet 2005) and determined if there were reductions in diversity estimates between 2003 and 2012 using Wilcoxon matched pairs rank sum tests (Zar 1996). We estimated the inbreeding coefficient (Fis) of each population in each sampling year using GenePop v 4.5.1 (Rousset 2008) to determine if there was evidence of inbreeding among populations and if this significantly differed according to sampling year. Finally, we examined the possibility that populations experienced genetic bottleneck using the program BOTTLENECK (Piry *et al*. 1999). This program examines the potential for greater expected heterozygosity based on allelic diversity relative to expected heterozygosity estimated under mutation-drift equilibrium (Nei *et al*. 1975; Cornuet & Luikart 1996). If a significantly high proportion of loci exhibit an allele deficiency relative to expectations based on mutation-drift equilibrium, the population would show signs of a recent reduction in the effective population size and thus a bottleneck (Nei *et al*. 1975; Cornuet & Luikart 1996). We conditioned analyses on the infinite alleles model (IAM), the step-wise mutation model (SMM) and the two-phase model (TPM) of microsatellite mutation since we are using microsatellites with a range of repeat motif types—dimeric, trimeric, and imperfect motifs—and thus we have no a priori reason to select one particular mutational model over another (repeat types presented in STable2 of (Kuester *et al*. 2015)). All analyses were performed across 1000 iterations assuming mutation-drift equilibrium, and significance was calculated using the Wilcoxon test (appropriate for sample sizes of < 30 individuals, (Luikart & Cornuet 1998; Luikart *et al*. 1998)).

**Resistance screen.** We examined the potential that populations and sampling years varied for resistance using univariate mixed-model analyses of variance. We assessed resistance in two ways—first, as a measure of the number of individuals within populations that died as a result of herbicide application, and second, as a measure of the amount of biomass change following herbicide application. We used the glmer option of the lme4 package in R (Bates *et al*. 2011) and modeled survival as a binary character (0/1) and used the lmer option to assess biomass remaining post-herbicide. In each model, replicate greenhouse experiment, herbicide treatment, collection year, and population were the independent variables with survival or standardized biomass as the dependent variables. We included interactions between population and collection year as well as population, collection year and treatment. Population and its interaction terms were considered random effects in each model whereas all other effects were fixed. We previously identified a significant population effect from the 2012 cohort for survival post-herbicide application, which indicated that populations vary in their respective level of resistance (Kuester *et al*. 2015). Here we are specifically interested in the year term as well as interaction terms including the year effect, which would indicate that resistance varies between sampling years and/or that populations vary in their level of resistance between years. An F-test was used to determine the significance of fixed effects, and the significance of each random effect in the model was determined using a likelihood ratio test (LRT) in which the full model was compared to a reduced model with the effect of interest removed. The *P*-value was determined using a |^2^ test with one degree of freedom. We examined the normality of our estimates of biomass with the Shapiro-Wilk test and by visual inspection of quantile-quantile (q-q) plot, and square root transformed this variable to improve normality of the residuals.

## Results

**Genetic diversity and differentiation**. We uncovered reductions in genetic diversity between sampling years among populations (Table 1), with most measures of diversity significantly reduced in 2012 compared to 2003 (Figure 2). For example, expected heterozygosity was 32% lower in 2012 (W = 51, P = 0.01), allelic richness was 18% lower (W = 52, P = 0.01), the effective number of alleles was 43% lower (W = 51, P = 0.01) and the absolute number of alleles per locus were reduced by 19% in 2012 compared to 2003 (W = 50, P = 0.01). The observed heterozygosity was 27% higher, on average, in 2012 compared to 2003 (W = 4, P = 0.005). This difference is likely due to the low observed compared to expected heterozygosity of the 2003 cohort, *i.e.,* the inbreeding coefficient (F_IS_ = 1 – H_o_/H_e_) was higher in 2003 versus 2012 (F_2003_ = 0.57 ± 0.05 (±SE) vs. F_2012_ = 0.13 ± 0.04, respectively; Figure 2). The difference in average F_IS_ value between 2003 and 2012 was significant (W = 55, P < 0.01). Although this difference could be due to selection against heterozygotes in 2003, it is more likely indicative of differences in the mating system between sampling years of this mixed-mating, hermaphroditic species. Populations were sampled during a slightly longer window of time in 2003 than in 2012 (10/10-11/3 in 2003 vs 10/15-10/20 in 2012); however, at least five of the 10 populations were sampled during the same temporal window (10/10-10/20 both years), and these populations exhibit similar differences in F values (F_2003_ = 0.47±0.08 vs. F_2012_ = 0.12±0.03). We do not have information regarding pollinator abundance or any other reason to expect differences in the mating system between years.

**Figure 2.**
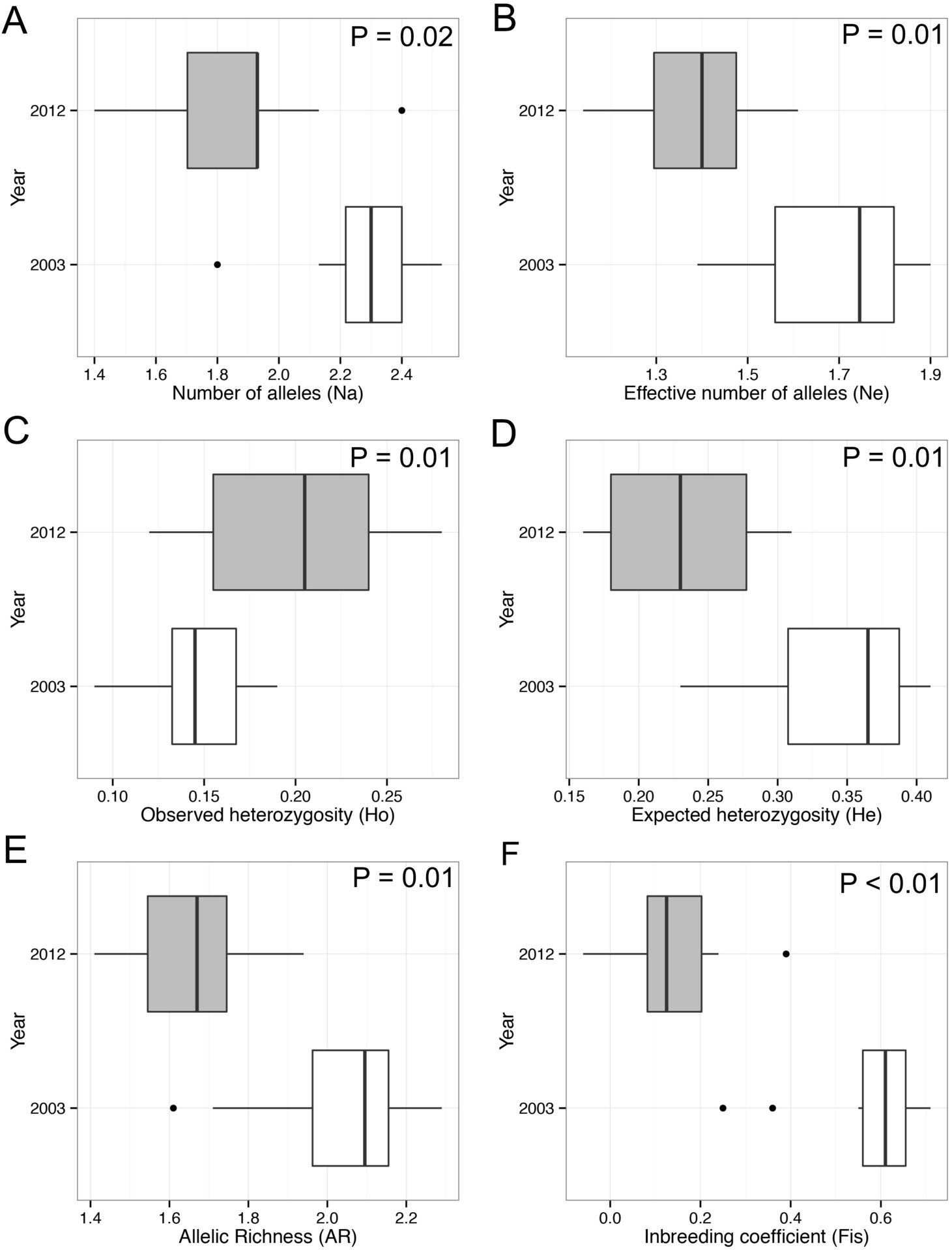
Genetic diversity indices compared between sampling years (2003 and 2012). Shown are the median (thick line) and lower and upper quartiles for (A) number of alleles (Na), (B) effective number of alleles (Ne), (C) observed heterozygosity (Ho), (D) expected heterozygosity (He), (E) allelic richness (AR), and (F) inbreeding coefficients (F_IS_).

**Table 1.** The genetic diversity of populations between sampling years. Shown are the number of alleles (Na), the effective number of alleles (Ne), the observed and expected heterozygosity (Ho and He, respectively), allelic richness (AR), and the inbreeding coefficient (F_IS_) of each population.

| Population | Na | | Ne | | Ho | | He | | AR | | $F_{IS}$ | |
| --- | --- | --- | --- | --- | --- | --- | --- | --- | --- | --- | --- | --- |
|  | 2003 | 2012 | 2003 | 2012 | 2003 | 2012 | 2003 | 2012 | 2003 | 2012 | 2003 | 2012 |
| 2 | 2.27 | 1.93 | 1.53 | 1.37 | 0.09 | 0.17 | 0.30 | 0.22 | 1.96 | 1.68 | 0.71 | 0.24 |
| 8 | 2.33 | 1.67 | 1.76 | 1.31 | 0.17 | 0.18 | 0.38 | 0.17 | 2.16 | 1.48 | 0.59 | -0.06 |
| 10 | 2.20 | 1.93 | 1.65 | 1.29 | 0.13 | 0.15 | 0.33 | 0.18 | 1.97 | 1.59 | 0.63 | 0.18 |
| 18 | 2.40 | 1.47 | 1.76 | 1.27 | 0.13 | 0.15 | 0.36 | 0.16 | 2.05 | 1.41 | 0.66 | 0.08 |
| 21 | 2.53 | 1.40 | 1.90 | 1.14 | 0.14 | 0.12 | 0.41 | 0.18 | 2.29 | 1.53 | 0.69 | 0.39 |
| 23 | 2.53 | 1.93 | 1.84 | 1.46 | 0.16 | 0.24 | 0.39 | 0.27 | 2.18 | 1.75 | 0.59 | 0.15 |
| 26 | 1.80 | 2.13 | 1.39 | 1.61 | 0.18 | 0.24 | 0.23 | 0.31 | 1.61 | 1.87 | 0.25 | 0.21 |
| 28 | 2.13 | 2.40 | 1.40 | 1.55 | 0.15 | 0.28 | 0.23 | 0.31 | 1.71 | 1.94 | 0.36 | 0.09 |
| 30 | 2.27 | 1.93 | 1.86 | 1.48 | 0.19 | 0.27 | 0.40 | 0.28 | 2.14 | 1.73 | 0.64 | 0.10 |
| 32 | 2.40 | 1.80 | 1.73 | 1.43 | 0.14 | 0.23 | 0.37 | 0.24 | 2.14 | 1.66 | 0.55 | 0.03 |

Bonferroni-corrected HWE tests, consequently, indicated that more loci were not in HWE equilibrium within populations in 2003 (39 out of 150 locus × population combinations), compared to 2012 (1 out of 150 locus × population combinations). Processes that lead to heterozygote deficit, such as inbreeding or population substructure can cause deviations from HWE; alternatively, the presence of null alleles could inflate estimates of homozygosity and lead to deviations from HWE (Kelly *et al*. 2011). We tested for the potential that null alleles altered our estimates of genetic diversity by removing loci with >25% putative null allele frequencies across populations and re-estimated indices of genetic diversity. We found no evidence that loci with potential null alleles altered our estimates of genetic diversity or the conclusion that diversity is altered between sampling years (Table S3, Supporting Information).

We further examined changes in the patterns of allelic diversity by investigating the number of alleles, the number of rare alleles (<10% frequency) and the frequency of rare alleles present in 2003 and 2012. At the species level (*i.e*., across all populations), we found no evidence for a reduction in the total number of alleles from 2003 to 2012 (42 versus 44 alleles in each year, respectively)—unexpectedly, we found fewer rare alleles in 2003 than 2012 (10 vs 17). Only four of the rare alleles present in 2003 were likewise present in 2012, and their frequency was not dramatically increased as would be expected if rare allele frequency changes were responsible for the higher observed heterozygosity in 2012. When examining the number of alleles per population, however, we found that the total number of alleles was reduced in eight of 10 populations, by as much as 12-40% across populations. Two of the ten populations (populations 26 and 28) exhibited gains of low frequency alleles (between 4-5 new alleles present in 2012 at frequencies of <10%). Thus, 8 of the 10 populations show reductions in diversity over time likely due to random genetic drift, whereas two of the populations exhibit an increase in the number of alleles, putatively due to migration, drift, or mutation.

We tested for a signature of bottleneck events in both the 2003 and 2012 samples using the BOTTLENECK program, and found that significance of these tests depended on both the specific model employed (IAM, SMM, or TPM) and the sampling year. Under the lAM, six populations sampled from 2003 exhibited significant heterozygote excess following corrections for multiple tests, whereas only one – population 32 – exhibited significant heterozygote excess under all three models of microsatellite evolution (Table 2). No populations from 2012 exhibited evidence of heterozygote excess and thus signs of a bottleneck following corrections for multiple tests (Table 2).

**Table 2.** Tests of heterozygosity excess within populations for each sampling year using the BOTTLENECK program (Cornuet & Luikart 1996). Tests were performed using three different models of microsatellite evolution, each of which assumes mutation-drift equilibrium (IAM, infinite alleles model; SMM, stepwise mutation model; TPM, two-phase model. Probability values from one-tailed Wilcoxon tests are shown, with bolded values indicating statistical significance following corrections for multiple tests (P < 0.005).

| Population | <u>2003</u> |  |  | <u>2012</u> |  |  |
| --- | --- | --- | --- | --- | --- | --- |
|  | IAM | SMM | TPM | IAM | SMM | TPM |
| 2 | 0.117 | 0.810 | 0.396 | 0.313 | 0.615 | 0.539 |
| 8 | **0.002** | 0.084 | 0.020 | 0.230 | 0.527 | 0.422 |
| 10 | 0.016 | 0.249 | 0.047 | 0.765 | 0.945 | 0.867 |
| 18 | **0.004** | 0.188 | 0.047 | 0.027 | 0.055 | 0.055 |
| 21 | **0.003** | 0.122 | 0.016 | 0.371 | 0.473 | 0.371 |
| 23 | **0.003** | 0.084 | 0.016 | 0.095 | 0.271 | 0.249 |
| 26 | 0.117 | 0.396 | 0.235 | 0.122 | 0.249 | 0.153 |
| 28 | 0.485 | 0.867 | 0.715 | 0.249 | 0.773 | 0.580 |
| 30 | **0.003** | 0.227 | 0.055 | 0.012 | 0.216 | 0.138 |
| 32 | **<0.001** | **0.004** | **0.001** | 0.032 | 0.170 | 0.133 |

We next estimated the effective number of individuals from each sampling year using expected heterozygosity and the equation H_e_ = 4N_e_*μ* (Nagylaki 1998) with a mutation rate, *μ*, of 10^−3^ (Marriage *et al*. 2009). We found that the estimated number of individuals from the 2003 populations was significantly higher, on average, compared to that of the 2012 populations (N_e, 2003_ = 85, N_e, 2012_ = 58; W = 67, P = 0.005). Furthermore, we found no significant difference between our census sample size from the 2012 populations and the estimated effective number of individuals from that sampling year (Population size average from census = 70 individuals; W = 36.5, P = 0.32). While the difference in estimated number of individuals between sampling years indicated that most populations experienced reductions in size (reductions ranging from 20-55 individuals fewer in 2012), populations 26 and 28 both exhibited an estimated gain of 20 individuals.

In line with lower diversity of the majority of populations, we found significant genetic differentiation between individuals sampled from different collection years (AMOVA year effect, F_RT_ = 0.218, P = 0.001, Table S4), and evidence that individuals sampled as seed in 2003 were more similar to one another than to individuals sampled as seed from the same location in 2012 (Figure 3), *i.e*., no individual assigned to 2003 was likewise assigned to 2012. We found that the estimate of F_RT_ was inflated by loci that potentially harbored null alleles; however, after removing these loci from analysis, we found that the F_RT_ estimate was still significantly different from zero, indicating the presence of between-year genetic differentiation (F_RT_(8 loci) = 0.133, P = 0.001, Table S4). We did not remove loci with potential null alleles from genotypic assignment as these tests are not greatly influenced by their presence (Carlsson 2008).

**Figure 3.**
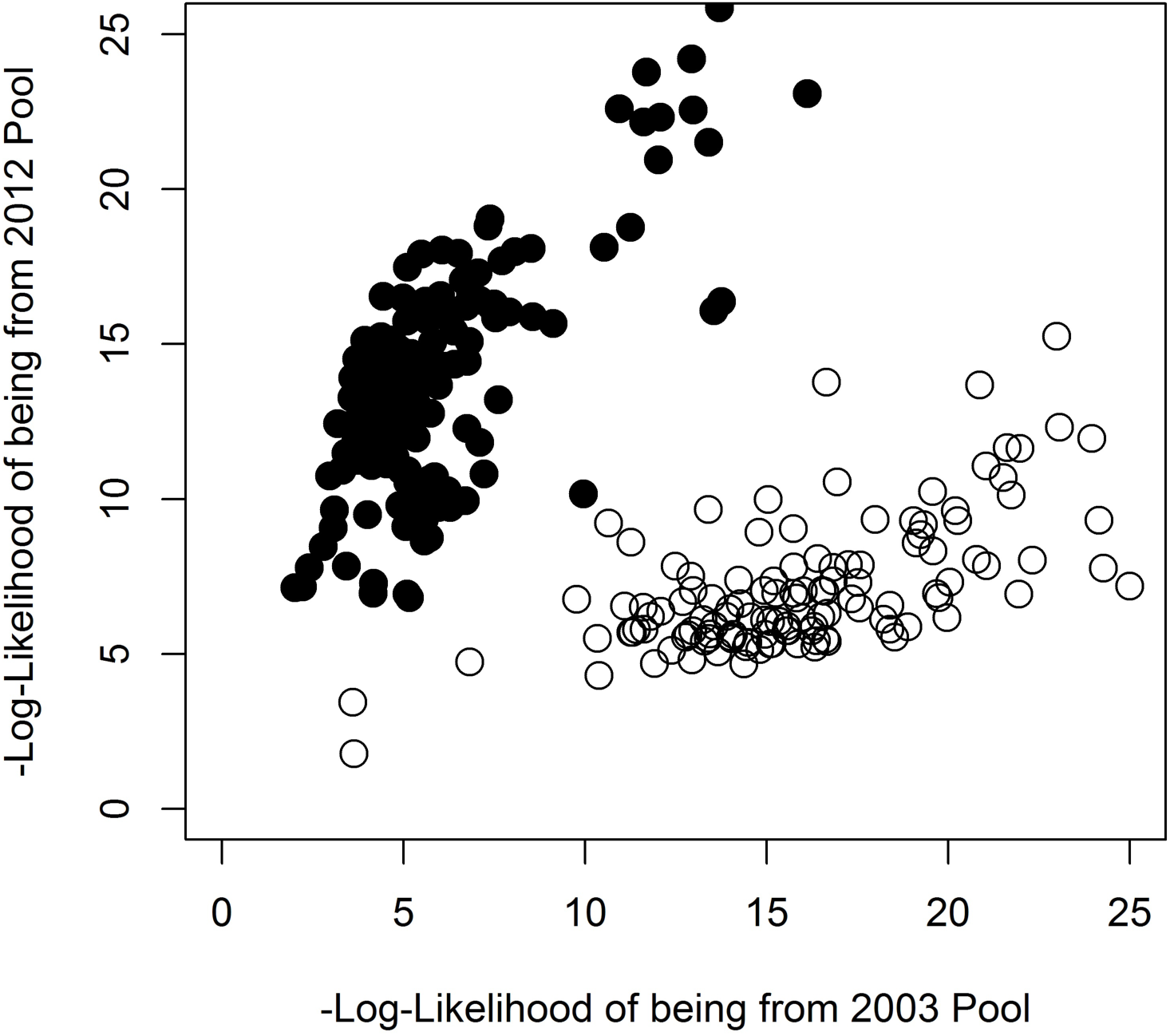
Scatter plots of log likelihood values from assignment tests of individual *I. purpurea* plants sampled in 2003 and 2012 based on genotypic data at 15 microsatellite loci. A higher position relative to the y-axis indicates a higher likelihood of being from 2012 pool of individuals and a higher position relative to the x-axis indicates greater likelihood of being from 2003 pool of individuals.

**Phenotypic divergence.** We examined resistance traits (survival and biomass post-herbicide application) to determine if there was evidence of changes in resistance between sampling years. Our mixed-effects analyses of variance uncovered a significant year effect for biomass remaining after herbicide application (F_1, 3595_ = 4.72, P = 0.03; Table 3). On average across all populations, the biomass remaining post-spray of the 2012 cohort was slightly greater than that of the 2003 cohort (62% vs. 57% in 2012 and 2003, respectively) suggesting moderate increases in resistance across populations sampled in 2012 (see Table S5 for averages (±SE) among all populations). Likewise, a higher percentage of individuals sampled in 2012 survived herbicide application compared to those sampled from 2003 (49% vs 42%), but this difference was not significant (F_1, 5365_ = 2.58, P = 0.11; Table 3).

**Table 3.** Generalized linear mixed effects model of resistance in *I. purpurea*. Models include fixed effects of experimental replicate, treatment, sampling year, sampling x treatment interaction; population and interactions of population x year, population x treatment, and population x treatment x year are considered random effects.

| Effect | Df | Survival |  | Df | Biomass |  |
| --- | --- | --- | --- | --- | --- | --- |
|  |  | F | P |  | F | P |
| <u>Fixed Effects</u> |  |  |  |  |  |  |
| Replicate | 1 | 14.46 | <0.001 | 1 | 12.58 | <0.001 |
| Treatment | 5 | 154.74 | <0.001 | 5 | 190.98 | <0.001 |
| Year | 1 | 2.58 | 0.108 | 1 | 4.72 | 0.030 |
| Year x Treatment | 5 | 0.09 | 0.994 | 5 | 0.86 | 0.511 |
| <u>Random Effects</u> |  | <u>X<sup>2</sup></u> |  |  | <u>X<sup>2</sup></u> |  |
| Population | 1 | 19.18 | <0.001 | 1 | 4.97 | 0.026 |
| Population x Year | 1 | 23.75 | <0.001 | 1 | 7.92 | 0.005 |
| Population x Treatment | 1 | 3.70 | 0.054 | 1 | <0.001 | 1 |
| Population x Treatment x Year | 1 | <0.001 | 1 | 1 | <0.001 | 1 |
| Residual DF | 5365 |  |  | 3595 |  |  |

As in previous work (Kuester *et al*. 2015), we identified significant population effects for both measures of resistance (Table 3), indicating that populations vary across the landscape for their relative level of herbicide resistance. Here, however, we also find population by year effects in each analysis, indicating that populations differ in their level of resistance across years (Survival, χ^2^ = 23.74, P < 0.001; Biomass, χ^2^ = 7.92, P = 0.005; Table 3). One population’s survival increased by 79% compared to 2003, indicating that some populations may respond more readily with increased resistance than others. In general, three populations sampled from TN that were highly resistant in 2012 (Kuester *et al*. 2015) were similarly resistant in 2003 (Figure 1 A & B, shown at 2X field rate of RoundUp). The majority of the significant increases identified in the 2012 cohort compared to the 2003 cohort were located in NC and SC (Figure 1 A & B)—while five populations from the 2012 cohort of this region exhibit resistance values significantly greater than the species-wide average (56% survival at 2X the field rate of RoundUp (Kuester *et al*. 2015)), the 2003 cohorts of these populations exhibited only ~14% survival at 2X field rate.

## Discussion

Despite the ubiquity and persistence of weedy plant populations, there are few examinations of how their neutral and adaptive genetic diversity may change over time. Here we use a resurrection ecology experiment to show that populations of weedy *I. purpurea* sampled from crop fields concomitantly lose genetic diversity and show signs of potential adaptive evolution in herbicide resistance. Our experiments yielded three novel findings. First, we found that seed progenies from populations sampled in 2012 exhibited lower genetic diversity and higher genetic differentiation than seed progenies sampled from the same fields and locations in 2003, suggesting that populations have experienced a genetic bottleneck between sampling periods. Second, heterozygosity excess tests indicated that a significant genetic bottleneck likely also occurred prior to 2003, perhaps due to the dramatic increase in glyphosate use in the late 1990’s (see Fig 1 in Baucom & Mauricio 2004). Although we cannot ascribe the loss of neutral genetic variation to the widespread use of herbicide *per se*, we show that a resistance trait—the amount of biomass maintained following herbicide application—has increased in some populations from 2003 to 2012. We combine these results with a retrospective analysis of a larger and previously published dataset showing that highly resistant populations sampled in 2012 exhibit significantly reduced heterozygosity and allelic richness estimates compared to less resistant populations. Below we discuss each of these major findings.

### Reductions in genetic diversity between sampling years

We currently have a very limited understanding of how agricultural regimes may influence the population genetics of agricultural weeds. Although processes such as tilling and herbicide use are hypothesized to result in genetic bottlenecks, previous examinations of agricultural weed populations have either failed to uncover substantial reductions in genomewide diversity (Kane & Rieseberg 2008) or have presented largely circumstantial evidence for bottlenecks (i.e., comparisons between species; (Hamrick *et al*. 1979)). The significant loss in diversity that we uncovered across populations of *I. purpurea* sampled from agricultural fields argues for a bottleneck that was either very strong or occurred more than once, or both. While there are no studies, to our knowledge, that examine the temporal genetics of agricultural weed populations for comparison, it is of note that the average loss of allelic richness that we identified across populations (on average 15% lower between cohorts) is similar in magnitude to that of introduced, colonizing species (18% loss compared to native populations (Dlugosch & Parker 2008)). Furthermore, using expected heterozygosity estimates from each sampling year, we find that the estimated population sizes have reduced between 2003 and 2012, with the majority of the populations losing reproductive individuals. While we did not take population census data in 2003 for comparison, we find that the estimated population size in 2012 is not significantly different from the census size, suggesting that our estimated population sizes are decent approximations of the true census size. The majority of the populations exhibited loss of alleles between sampling years, however, two populations—#26 and #28—exhibited gains of low frequency alleles, and the estimated sample size of these two populations likewise increased relative to other populations. The increased diversity of these populations likely were from dormant seeds, or, alternatively, due to migration from another population. Emergence of seed stored in the seed bank is incredibly likely—this species can produce a large number of seeds in field conditions (between 3,000-10,000 per individual), and these heavy, gravity dispersed seeds can remain dormant for ~20 years in the soil (Baskin & Baskin 2000).

Interestingly, while the loss of allelic diversity between 2003 and 2012 suggests a genetic bottleneck has occurred between sampling years, the within-population examination of heterozygosity excess (i.e, the bottleneck test) did not find evidence of genetic bottleneck in the 2012 populations. Instead, our tests of heterozygosity excess uncovered evidence of genetic bottleneck among six populations collected in 2003 under the IAM model, with one population exhibiting evidence of a genetic bottleneck under all three models of microsatellite evolution. Other studies using heterozygosity excess tests have reported limited support of genetic bottleneck in species known to have experienced population declines (Hufbauer *et al*. 2004; Peery *et al*. 2012) with the general conclusion that heterozygosity-excess tests may be limited to severely bottlenecked populations (Peery *et al*. 2012). Our results suggest that a genetic bottleneck occurred in some populations prior to the 2003 sampling, possibly following the sharp increase in the widespread use of glyphosate across RoundUp Ready crops (see Fig 1 in Baucom & Mauricio 2004). In this scenario, the loss of allelic diversity that we identified in 2012 may simply be a continuation of effects following the initial population bottleneck. Alternatively, it is possible that demographic bottlenecks occurred prior to 2003 and continued between 2003 and 2012, but to a lesser extent between years. While a few of the 2012 populations did exhibit heterozygosity excess, defined as greater heterozygosity based on estimates of allelic diversity relative to heterozygosity estimated under mutation-drift equilibrium, tests were not significant following corrections.

Another attribute of the data suggest populations experienced a genetic bottleneck prior to 2003—we found more locus by population combinations out of HWE compared to 2012. In addition to the evolutionary forces of genetic drift, selection, mutation, migration, and nonrandom mating (i.e. inbreeding or non-assortative mating), null alleles can also cause deviations from Hardy-Weinberg equilibrium and will appear as heterozygote deficiency (Dąbrowski *et al*. 2014). The pattern that we uncovered of low observed heterozygosity relative to expected heterozygosity in our 2003 populations is consistent with the presence of null alleles but also consistent with high levels of inbreeding. To investigate the potential that null alleles influenced our results, we removed loci with putative null alleles from analyses and found that estimates of genetic diversity remained lower in the 2012 populations compared to their 2003 counterparts; further, F indices remained significantly higher in the 2003 sample, implicating wide-spread inbreeding following a potential bottleneck. *I. purpurea* is a hermaphroditic species that displays a wide range of outcrossing rates in nature (*t*_m_ range: 0.2-0.8, across 20 populations; Kuester et al. *unpublished data*), thus it is plausible that inbreeding could follow a large demographic bottleneck in this species. The majority of our loci exhibited F_IS_ > 0 in 2003 across populations, further pointing to a scenario of inbreeding following a large demographic change rather than the influence of null alleles. Finally, it is of note that null allele detection methods have been shown to exhibit low reliability when applied to non-equilibrium populations and will overestimate their frequency when populations have recently experienced demographic bottleneck (Dąbrowski *et al*. 2014).

### Pattern of phenotypic variation between temporal samples

Recent work provides an interesting contrast between the phenotypic and neutral genetic variation spatially distributed within this system—while neutral genetic differentiation among 44 *I. purpurea* populations is low (*i.e*., F_ST_ = 0.127), populations are significantly differentiated for herbicide resistance across the landscape, with some populations exhibiting high and others very low resistance (Kuester *et al*. 2015). Our screen of herbicide resistance in 26 of these temporally sampled populations shows that, in addition to a mosaic of resistance across the landscape, the level of resistance has slightly increased, on average, between sampling dates. This finding is interesting in light of the reduced neutral genetic variation that we identified in eight of our 10 temporally sampled populations, and alternatively, in light of evidence of potential migrants in two of the 10 populations—reductions in diversity as well as influx of presumably non-adapted variation would either act to impede or to counteract adaptation. These forces, along with recent work showing a severe fitness penalty of herbicide resistance in this species (Van Etten *et al*. 2015) likely explain why the average increase in resistance that we identified among all populations was modest—perhaps the populations that maintained resistance between sampling years (TN populations) or those that exhibit large increases in resistance (NC/SC populations) were less influenced by susceptible migrants germinating from the seed bank and/or costs of resistance than other populations. The low average increase in resistance across populations could also be due to a range of other factors: it is possible that few populations house additive genetic diversity for resistance, populations may experience different selective regimes, or the response to selection via glyphosate has been constrained by bottleneck events. Interestingly, there was no evidence that plants were different in size between the years (*data not shown*), indicating that the increased resistance we detected is not due to plants simply being larger in the 2012 cohort and thus better able to withstand herbicide application.

Although we find evidence for a moderate increase in the level of resistance across populations, it is important to note that our phenotypic comparisons were made using field-collected seeds. Thus our resistance estimates include both potential genetic components and environmental effects. This could explain the slight increase in resistance over time: if more 2012 populations had experienced glyphosate application relative to the 2003 populations, we would perhaps be sampling from a subset of the population that experienced herbicide relative to plants that had not, potentially inflating estimates of resistance in the 2012 samples. In 2003, between 80-92% of the soy fields in the US were RoundUp Ready, and thus sprayed with the herbicide, whereas approximately 20% of corn was RoundUp Ready in that sampling year. In 2012, 93% of soy planted was RoundUp Ready and between 73-80% of corn was RoundUp Ready (NASS 2015). Thus, it is possible that our comparison of the temporally sampled phenotypes is influenced by exposure to the herbicide itself in 2012. We considered this by using crop type as a proxy for herbicide use, and determined if biomass remaining post-spray differed according to crop (restricted to soy and corn) separately for both 2003 and 2012. There was no difference in biomass remaining post-herbicide according to crop either year—e.g., the biomass remaining post-herbicide of morning glories sampled from soy was no different than those sampled from corn—suggesting that the population-level estimates of resistance are not dependent on the crop type (as a proxy for spraying regime) in one particular sampling year. While we cannot conclude that the increase in our field estimates of resistance between sampling years is solely due to adaptive evolution, we have previously identified an additive genetic basis underlying glyphosate resistance in one population of this species (Baucom & Mauricio 2008), positive selection for increased resistance in the field (Baucom & Mauricio 2008), and further, have shown that resistance segregates in crosses (Debban *et al*. 2015) indicating that the genetic potential is present within at least one population. Identification of the genetic basis of resistance across populations, and an assessment of how alleles associated with resistance change over time will decisively test our hypothesis that selection from the use of this herbicide is leading to adaptation in natural populations.

### Has herbicide application caused the genetic bottleneck?

The populations used in this study were all sampled from crop fields that were farmed prior to and from 2003 onward. While we do not have specific information on herbicide use over this time period, we have historical record for six of 10 years (Table S1) showing that these locations were used for corn and soy crops, both of which make use of herbicides—and largely glyphosate—for weed control. Although other environmental factors (e.g., those associated with climate change) could be responsible for the genetic bottleneck that we report herein, herbicide use is an obvious potential factor. We examined this idea by making use of a larger and previously published dataset of 32 populations, sampled in 2012, for which we have estimates of both resistance and genetic diversity (Kuester *et al*. 2015). We performed separate regressions of expected heterozygosity (He) and allelic richness (AR) on estimates of herbicide resistance (proportion survival) at 3.4 kg ai/ha of glyphosate and found a significant negative relationship, *i.e*., more resistant populations exhibit lower genetic diversity (He vs resistance: R = -0.375, P = 0.04; AR versus resistance: R = -0.345, P = 0.05). There is thus some indication that selection via herbicide application has led to the genetic bottleneck among populations. Interestingly, a population used in both analyses – population 32 – exhibits high survival at 3.4 kg ai/ha of glyphosate, and was the population for which we found evidence of a pre-2003 bottleneck under all models of microsatellite evolution.

Conclusions—Weedy plant species found in agricultural fields experience strong selection and thus are hypothesized to be either plastic, capable of adaptation, or saved from extinction through gene flow (Baker 1974; Parker *et al*. 2003). By using a resurrection ecology framework, we provide evidence that even though genetic variation is lost from the system, some populations show potential signs of adaptation to herbicide application. While previous work indicates that the majority of the gene flow across southeastern populations occurred prior to the wide-spread adoption and use of glyphosate, suggesting that resistance evolution is due to selection on standing or novel variation within populations, that we identified evidence of potential migrants into the 2012 gene pool (in at least two populations) does not allow us to rule out the hypothesis that resistance can be introduced from outside sources.

Further, while we find evidence of increased resistance, we also show that the absolute change between years was not drastic; large resistance gains were limited to particular populations. These data suggest heightened measures should be taken to reduce the likelihood that seeds are accidentally moved between crop fields through farm machinery or through contaminated seed lots. Finally, we have some evidence that the lower genome-wide diversity identified across populations is due to the application of glyphosate; however, we note that we cannot rule out other potential factors, since other herbicides with different mechanisms of action are often applied in crops, other cropping techniques that reduce population sizes might have been employed, and it is also possible that populations have lost diversity due to changes in the climate. The results shown here suggest that this weed, while being a ‘general purpose genotype’ (Baker 1974; Chaney & Baucom 2014) is also capable of adaptive evolution even while losing significant allelic diversity. How likely future adaptation to novel selective forces may be in the future, in light of reduced variation, is unknown.

## Acknowledgements

The authors wish to thank M. Van Etten, Y. Brandvain, D. Alvarado Serrano, and S. Colom for providing feedback that improved this manuscript, and A. Wilson, E. Fall, A. Teodorescu, S. Colom, and S. Smitka for assistance with greenhouse and lab work. This work was funded by USDA NIFA grants 04180 and 07191 to RSB and SMC.

## Data Accessibility

SSR Genotyping and morphological data is available on Dryad, doi to be provided upon acceptance.

Sampling locations and crop history information is provided in the Supplemental materials section.

## Supporting Information

### Materials and Methods

*Scoring accuracy and allelic composition of populations/sampling years*—We estimated the scoring error rates per locus by randomly selecting and re-scoring the genotypes of 200 individuals. We found our scoring error rate to be very low across loci: IP31 (0%), IP2 (3.7%), IP27 (0%), IP8 (0%), IP34 (0%), IP1 (0%), IP36 (6.52%), IP47 (1.45%), IP12 (1.45%), IP21 (5.07%), IP6 (1.45%), IP45 (2.17%), IP26 (2.17%), IP42 (3.62%).

The total number of alleles observed per locus within 184 individuals sampled in 2003 and 171 individuals sampled in 2012 samples ranged from 2 to 7 in 2003 and 1 to 7 in 2012. Overall, we found a total of 42 alleles scored across 15 loci in 2003 and 44 alleles in 2012. On average there were 2.3 alleles per locus x population combination in 2003 and 1.9 alleles in 2012. The size range of alleles ranged roughly between 90 and 300 base pairs.

*Hardy-Weinberg Equilibrium, scoring errors, and null alleles*—Deviations from Hardy-Weinberg equilibrium (HWE) were tested using GenePop v 4.5.1 (Rousset 2008). After performing Bonferroni corrections (Rice 1989) for multiple tests, we found 39 locus x population combinations that departed from HWE in the 2003 and 1 locus x population combination that deviated from HWE in the 2012 data, respectively, out of 150 possible locus x population combinations. In 2003, the number of loci out of HWE per population ranged from 0-8 (median: 2); the departures in HWE among loci were not consistent across populations. Small population size, non-random mating, mutations, migration, and selection can cause departures in HWE; furthermore, null alleles can lead to heterozygote deficiency, which may be detected as departures from HWE. Therefore, we further explored the data to determine if null alleles and other genotyping errors were responsible for the higher number of loci out of HWE among the 2003 compared to 2012 populations.

We assessed scoring errors per locus x population combination using MicroChecker v 2.2.3 (Van Oosterhout et al. 2004) and GenePop v 4.5.1 (Rousset 2008). Large allele dropout (underamplification of larger allelic variants) and scoring errors due to stutter (an error resulting from slippage of the polymerase during PCR) were not detected in our dataset. MicroChecker v 2.2.3 (Van Oosterhout et al. 2004) identified more loci with potential null alleles in the 2003 data compared to the 2012 data (8 of 15 versus 3 of 15 loci, respectively). We examined the frequencies of potential null alleles across loci × population combinations using GenePop v 4.5.1 and identified loci (IP2f, IP27f, IP18f, IP12f, IP21f, IP26f, IP42f) with null alleles at a frequency of >25%, on average, across populations. We removed these loci from both the 2003 and 2012 datasets (leaving eight loci: IP31, IP8, IP34f, IP1f, IP36, IP47f, IP6f, IP45f) and estimated expected heterozygosity, allelic richness, effective and absolute number of alleles using GenAlEx v. 6.5 (Pekall and Smouse 2012). All diversity estimates remained significantly lower in the 2012 data (Table S3) indicating that null alleles were not responsible for the reduced genetic variation in the 2012 populations compared to the 2003 populations. We likewise estimated genetic differentiation between years using hierarchical AMOVA in GenAlEx v. 6.5 (Pekall and Smouse 2012) and found that, as expected based on previous reports (Carlsson 2008), our F_RT_, or measure of between-year differentiation, was potentially inflated by the seven loci with potential null alleles (Table S4, A and B: 15 loci F_RT_ = 0.218; 8 loci F_RT_ = 0.133). We elected to present results using all 15 loci in this work given that our patterns of diversity loss remained when these seven loci were removed from analysis, and in light of reports showing that bottleneck events lead to false-positives in null allele detection methodologies (Dabrowski et al. 2014). We did not remove loci with potential null alleles when performing assignment testing since null alleles do not significantly alter the outcome of such assignments (Carlsson 2008).

*Linkage disequilibrium*—We tested the presence of linkage disequilibrium between locus pairs for each population x sampling year combination using GenePop v 4.5.1 (Rousset 2008). All results were adjusted for multiple comparisons using Bonferroni correction (Rice 1989). Of 105 tested paired locus tests over either 2003 or 2012 sampled populations for linkage disequilibrium, 36 were found significant in 2003, though no disequilibrium was uncovered in 2012 (P < 0.05). The majority of locus pairs in disequilibrium from 2003 were from population 26 (25 significant pairs).

### Supplementary Tables

**Table S1.** Site and location information for *I. purpurea* populations used in genetic, growth and herbicide resistant assays. Shown are the population number, state of each population, the crop present in the field from 2008 to 2012, along with Latitude and Longitude GPS coordinates.

| Population Number | State | Field Type |  |  |  |  |  | Latitude | Longitude |
| --- | --- | --- | --- | --- | --- | --- | --- | --- | --- |
|  |  | 2003 | 2008 | 2009 | 2010 | 2011 | 2012 |  |  |
| 2 | NC | soy | corn | soy | corn | soy | corn | 34.595714 | -77.927484 |
| 4 | NC | soy | corn | soy | corn | soy | corn | 34.556672 | -79.125602 |
| 9 | NC | soy | na | soy | soy | soy | soy | 34.924044 | -77.796171 |
| 10 | NC | soy | corn | corn | soy | corn | soy | 34.983161 | -78.039309 |
| 11 | NC | soy | corn | soy | corn | soy | corn | 34.527135 | -78.756704 |
| 14 | NC | na | cotton | soy | soy | soy | soy | 35.424763 | -77.917121 |
| 19 | NC | soy | corn | soy | corn | tobacco | corn | 34.508193 | -78.70899 |
| 21 | NC | soy | cotton | soy | corn | cotton | soy | 35.369816 | -77.877314 |
| 22 | NC | soy | soy | cotton | corn | corn | corn | 36.1436 | -78.053422 |
| 25 | NC | soy | soy | soy | fallow | soy | corn | 34.616361 | -79.051667 |
| 29 | NC | soy | corn | soy | corn | soy | corn | 34.705135 | -78.738897 |
| 5 | SC | soy | soy | soy | soy | corn | soy | 33.859875 | -79.909072 |
| 8 | SC | na | corn | soy | corn | soy | corn | 34.297195 | -79.991259 |
| 12 | SC | soy | soy | corn | soy | cotton | Cotton | 34.145812 | -79.865313 |
| 15 | SC | soy | soy | soy | peanut | soy | soy | 34.104209 | -79.073735 |
| 16 | SC | soy | fallow | fallow | fallow | peanut | Alfalfa | 34.10535 | -79.183234 |
| 17 | SC | soy | soy | soy | soy | soy | soy | 34.159155 | -79.272908 |
| 18 | SC | soy | soy | soy | corn | soy | corn | 34.156593 | -79.27027 |
| 28 | SC | soy | cotton | corn | soy | cotton | corn | 34.097917 | -80.377715 |
| 47 | SC | na | soy | corn | soy | cotton | soy | 34.282132 | -79.746597 |
| 1 | TN | corn | alfalfa | alfalfa | alfalfa | soy | corn | 35.775237 | -85.903419 |
| 20 | TN | soy | soy | soy | soy | soy | corn | 35.830692 | -85.777871 |
| 23 | TN | soy | corn | soy | corn | soy | corn | 35.067905 | -86.62955 |
| 26 | TN | soy | corn | soy | soy | corn | soy | 35.533413 | -85.951902 |
| 30 | TN | corn | corn | soy | corn | soy | corn | 35.31105 | -85.945003 |
| 31 | TN | corn | soy | corn | soy | soy | corn | 35.608482 | -85.846379 |
| 32 | TN | soy | soy | soy | soy | soy | corn | 35.099356 | -86.225509 |

**Table S2.** Sampling information for temporally-sampled populations of *Ipomoea purpurea*. Shown are the population identification numbers and sample sizes by year (2003 or 2012) for the SSR genotyping and resistance assays.

| Population Number | SSR Sample Size |  | Resistance Assay Sample Size |  |
| --- | --- | --- | --- | --- |
|  | 2003 | 2012 | 2003 | 2012 |
| 2 | 16 | 15 | 113 | 101 |
| 4 | -- | -- | 117 | 115 |
| 9 | -- | -- | 113 | 87 |
| 10 | 17 | 26 | 115 | 68 |
| 11 | -- | -- | 90 | 116 |
| 14 | -- | -- | 115 | 116 |
| 19 | -- | -- | 113 | 106 |
| 21 | 16 | 6 | 116 | 109 |
| 22 | -- | -- | 114 | 107 |
| 25 | -- | -- | 116 | 100 |
| 29 | -- | -- | 113 | 112 |
| 5 | -- | -- | 118 | 69 |
| 8 | 13 | 20 | 110 | 107 |
| 12 | -- | -- | 116 | 117 |
| 15 | -- | -- | 105 | 118 |
| 16 | -- | -- | 105 | 112 |
| 17 | -- | -- | 114 | 118 |
| 18 | 22 | 15 | 115 | 115 |
| 28 | 27 | 18 | 109 | 117 |
| 1 | -- | -- | 63 | 62 |
| 20 | -- | -- | 59 | 61 |
| 23 | 24 | 17 | 80 | 110 |
| 26 | 18 | 16 | 108 | 115 |
| 30 | 12 | 20 | 115 | 115 |
| 31 | -- | -- | 110 | 105 |
| 32 | 19 | 18 | 66 | 75 |

**Table S3.** The genetic diversity of populations between sampling years with loci with putative null alleles removed (retaining 8 of the original 15). Shown are the number of alleles (Na), the effective number of alleles (N_e_), the observed and expected heterozygosity (Ho and He, respectively), and the fixation indices estimated by GeneAlEx v 6.5 (Pekall and Smouse 2012).

|  | Na |  | Ne |  | Ho |  | He |  | F |  |
| --- | --- | --- | --- | --- | --- | --- | --- | --- | --- | --- |
| Population | 2003 | 2012 | 2003 | 2012 | 2003 | 2012 | 2003 | 2012 | 2003 | 2012 |
| 2 | 2.286 | 2.143 | 1.634 | 1.386 | 0.161 | 0.267 | 0.338 | 0.224 | 0.478 | -0.145 |
| 8 | 2.571 | 1.714 | 1.892 | 1.424 | 0.221 | 0.328 | 0.421 | 0.211 | 0.454 | -0.419 |
| 10 | 2.143 | 2.000 | 1.529 | 1.313 | 0.218 | 0.225 | 0.295 | 0.198 | 0.429 | -0.108 |
| 18 | 2.714 | 1.571 | 1.840 | 1.439 | 0.214 | 0.267 | 0.364 | 0.246 | 0.463 | -0.041 |
| 21 | 2.857 | 1.714 | 2.112 | 1.305 | 0.179 | 0.262 | 0.451 | 0.212 | 0.523 | -0.227 |
| 23 | 2.857 | 2.000 | 1.970 | 1.551 | 0.269 | 0.353 | 0.432 | 0.308 | 0.367 | -0.056 |
| 26 | 2.000 | 2.143 | 1.530 | 1.488 | 0.246 | 0.366 | 0.316 | 0.269 | 0.207 | -0.203 |
| 28 | 2.286 | 2.714 | 1.548 | 1.504 | 0.231 | 0.413 | 0.319 | 0.303 | 0.157 | -0.131 |
| 30 | 2.571 | 1.857 | 1.774 | 1.492 | 0.173 | 0.379 | 0.374 | 0.272 | 0.451 | -0.167 |
| 32 | 2.429 | 2.143 | 1.864 | 1.467 | 0.250 | 0.379 | 0.376 | 0.274 | 0.425 | -0.143 |
| Average | 2.471 | 2.000 | 1.769 | 1.437 | 0.216 | 0.324 | 0.369 | 0.252 | 0.395 | -0.164 |
| Wilcoxon<br>paired<br>rank sums | <0.01 |  | <0.01 |  | <0.01 |  | <0.01 |  | <0.01 |  |

**Table S4.** Analysis of Molecular Variance (AMOVA) of neutral genetic data estimated using GeneAlEx v 6.5 (Pekall and Smouse 2012). Shown separately for A) 15 and B) 8 loci are the main effects of sampling year (2003 vs 2012), population nested within year and individuals nested within populations, and F and P values for each effect.

| **A.** |  |  |  |  |
| --- | --- | --- | --- | --- |
| Source | df | F-Statistics | Value | P |
| Year | 1 | Frt | 0.218 | 0.001 |
| Population(Year) | 18 | Fsr | 0.179 | 0.001 |
| Individual (Population) | 335 | Fst | 0.358 | 0.001 |
| Individual (Total) | 355 | Fis | 0.522 | 0.001 |
| Total | 709 | Fit | 0.693 | 0.001 |

| **B.** |  |  |  |  |
| --- | --- | --- | --- | --- |
| Source | df | F-Statistics | Value | P |
| Year | 1 | Frt | 0.133 | 0.001 |
| Population(Year) | 18 | Fsr | 0.167 | 0.001 |
| Individual (Population) | 335 | Fst | 0.278 | 0.001 |
| Individual (Total) | 355 | Fis | 0.320 | 0.001 |
| Total | 709 | Fit | 0.509 | 0.001 |

**Table S5.** Mean and standard error for (A) resistance traits (survival and above-ground biomass at 1.7 kg a.i./ha glyphosate) and (B) size traits (plant height and number of leaves) for each population x year combination. Survival was calculated as the number of individuals per population that survived glyphosate application divided by the total number of individuals within the population. Populations 1, 20 and 32 were removed from analysis of biomass remaining post-herbicide due to very low sample size (range: 0-6 per treatment).

| PopID | Survival |  | Biomass |  |  |  |
| --- | --- | --- | --- | --- | --- | --- |
|  | 2003 | 2012 | 2003 |  | 2012 |  |
|  | Proportion | Proportion | Mean | SE | Mean | SE |
| 1 | 1.000 | 1.000 | -- |  | -- |  |
| 2 | 0.278 | 0.353 | 0.575 | 0.120 | 0.395 | 0.051 |
| 4 | 0.158 | 0.211 | 0.430 | 0.035 | 0.466 | 0.058 |
| 5 | 0.263 | 0.727 | 0.479 | 0.051 | 0.449 | 0.075 |
| 8 | 0.647 | 0.529 | 0.755 | 0.130 | 0.688 | 0.130 |
| 9 | 0.474 | 0.538 | 0.364 | 0.043 | 0.513 | 0.089 |
| 10 | 0.211 | 1.000 | 0.436 | 0.052 | 0.328 | 0.113 |
| 11 | 0.400 | 0.471 | 0.371 | 0.073 | 0.618 | 0.099 |
| 12 | 0.105 | 0.200 | 0.453 | 0.056 | 0.521 | 0.051 |
| 14 | 0.368 | 0.150 | 0.552 | 0.066 | 0.457 | 0.088 |
| 15 | 0.444 | 0.263 | 0.564 | 0.099 | 0.602 | 0.095 |
| 16 | 0.353 | 0.263 | 0.266 | 0.034 | 0.429 | 0.055 |
| 17 | 0.150 | 0.250 | 0.392 | 0.029 | 0.517 | 0.053 |
| 18 | 0.211 | 0.389 | 0.450 | 0.080 | 0.479 | 0.041 |
| 19 | 0.333 | 0.667 | 0.383 | 0.046 | 0.524 | 0.049 |
| 20 | 0.900 | 1.000 | -- |  | -- |  |
| 21 | 0.368 | 0.438 | 0.545 | 0.089 | 0.508 | 0.055 |
| 22 | 0.667 | 0.421 | 0.406 | 0.059 | 0.502 | 0.120 |
| 23 | 0.615 | 0.500 | 0.284 | 0.106 | 0.536 | 0.095 |
| 25 | 0.368 | 0.313 | 0.427 | 0.044 | 0.290 | 0.031 |
| 26 | 0.421 | 0.368 | 0.424 | 0.080 | 0.498 | 0.057 |
| 28 | 0.375 | 0.316 | 0.398 | 0.059 | 0.381 | 0.035 |
| 29 | 0.316 | 0.611 | 0.459 | 0.043 | 0.601 | 0.073 |
| 30 | 0.350 | 0.421 | 0.417 | 0.044 | 0.376 | 0.060 |
| 31 | 0.235 | 0.400 | 0.396 | 0.073 | 0.422 | 0.064 |
| 32 | 0.900 | 0.833 | -- |  | -- |  |

## References

Baker HG (1974) The evolution of weeds. Annual Review of Ecology and Systematics, 5, 1–25.

Barrett SCH (1983) Crop mimicry in weeds. Economic Botany, 37, 255–282.

Barrett SCH (1988) Genetics and evolution of agricultural weeds. In: Weed Management in Agroecosystems Ecological Approaches (eds Altieri M, Liebman MZ), pp. 57–75. CRC Press, Boca Raton.

Baskin CC, Baskin JM (2000) Seeds. Academic Press, San Diego.

Bates D, Maechler M, Bolker B (2011) lme4: linear mixed-effects models using S4 classes.

Baucom RS, Chang S-M, Kniskern JM, Rausher MD, Stinchcombe JR (2011) Morning glory as a powerful model in ecological genomics: tracing adaptation through both natural and artificial selection. Heredity, 107, 377–385.

Baucom RS, Mauricio R (2004) Fitness costs and benefits of novel herbicide tolerance in a noxious weed. Proceedings of the National Academy of Sciences of the United States of America, 101, 13386–13390.

Baucom RS, Mauricio R (2008) Constraints on the evolution of tolerance to herbicide in the common morning glory: resistance and tolerance are mutually exclusive. Evolution, 62, 2842–2854.

Baucom RS, Mauricio R (2010) Defence against the herbicide Round Up (R) predates its widespread use. Evolutionary Ecology Research, 12, 131–141.

Baudouin L, Lebrun P (2000) An operational Bayesian approach for the identification of sexually reproduced cross fertilized populations using molecular markers. In: International Symposium on Molecular Markers for Characterizing Genotypes and Identifying Cultivars in Horticulture 546, pp. 81–93.

Carlsson J (2008) Effects of microsatellite null alleles on assignment testing. The Journal of Heredity, 99, 616–623.

Chaney L, Baucom RS (2014) The costs and benefits of tolerance to competition in *Ipomoea purpurea*, the common morning glory. Evolution, 68, 1698–1709.

Clegg MT, Durbin ML (2000) Flower color variation: A model for the experimental study of evolution. Proceedings of the National Academy of Sciences, 97, 7016–7023.

Cornuet JM, Luikart G (1996) Description and power analysis of two tests for detecting recent population bottlenecks from allele frequency data. Genetics, 144, 2001–2014.

Cornuet JM, Piry S, Luikart G, Estoup A, Solignac M (1999) New methods employing multilocus genotypes to select or exclude populations as origins of individuals. Genetics, 153, 1989–2000.

Dąbrowski MJ, Pilot M, Kruczyk M et al. (2014) Reliability assessment of null allele detection: inconsistencies between and within different methods. Molecular Ecology Resources, 14, 361–373.

Debban CL, Okum S, Pieper KE, Wilson A, Baucom RS (2015) An examination of fitness costs of glyphosate resistance in the common morning glory, Ipomoea purpurea. Ecology and Evolution, 5, 5284–5294.

Defelice MS (2001) Tall morningglory, *Ipomoea purpurea* (L.) Roth - Flower or Foe? Weed technology: a journal of the Weed Science Society of America, 15, 601–606.

Délye C, Jasieniuk M, Le Corre V (2013) Deciphering the evolution of herbicide resistance in weeds. Trends in Genetics: TIG, 29, 649–658.

Dlugosch KM, Parker IM (2008) Founding events in species invasions: genetic variation, adaptive evolution, and the role of multiple introductions. Molecular Ecology, 17, 431–449.

Fang Z, Gonzales AM, Durbin ML et al. (2013) Tracing the geographic origins of weedy *Ipomoea purpurea* in the southeastern United States. The Journal of Heredity, 104, 666–677.

Franks SJ, Sim S, Weis AE (2007) Rapid evolution of flowering time by an annual plant in response to a climate fluctuation. Proceedings of the National Academy of Sciences, 104, 1278–1282.

Goudet J (2005) FSTAT (Version 1.2): A Computer Program to Calculate F-Statistics. The Journal of Heredity, 86, 485–486.

Hamrick JL, Linhart YB, Mitton JB (1979) Relationships between life history characteristics and electrophoretically detectable genetic variation in plants. Annual Review of Ecology and Systematics, 10, 173–200.

Heap, I (2016) International survey of herbicide resistant weeds. Online. Internet. Tuesday, March 29, 2016. Available www.weedscience.com

Hufbauer RA, Bogdanowicz SM, Harrison RG (2004) The population genetics of a biological control introduction: mitochondrial DNA and microsatellite variation in native and introduced populations of *Aphidus ervi*, a parisitoid wasp. Molecular Ecology, 13, 337–348.

Jasieniuk M, BruleBabel AL, Morrison IN (1996) The evolution and genetics of herbicide resistance in weeds. Weed Science, 44, 176–193.

Kane NC, Rieseberg LH (2008) Genetics and evolution of weedy *Helianthus annuus* populations: adaptation of an agricultural weed. Molecular Ecology, 17, 384–394.

Kelly AC, Mateus-Pinilla NE, Douglas M et al. (2011) Microsatellites behaving badly: empirical evaluation of genotyping errors and subsequent impacts on population studies. Genetics and Molecular Research: GMR, 10, 2534–2553.

Kuester A, Baucom RS, Chang S-M (2015) The geographic mosaic of herbicide resistance evolution in the common morning glory, *Ipomoea purpurea*: Evidence for resistance hotspots and low genetic differentiation across the landscape. Evolutionary Applications, In Press, 1–44.

Lenski RE (1998) Bacterial evolution and the cost of antibiotic resistance. International microbiology: the official journal of the Spanish Society for Microbiology, 1, 265–270.

Lenski RE, Travisano M (1994) Dynamics of adaptation and diversification: a 10,000-generation experiment with bacterial populations. Proceedings of the National Academy of Sciences, 91, 6808–6814.

Luikart G, Allendorf FW, Cornuet JM, Sherwin WB (1998) Distortion of allele frequency distributions provides a test for recent population bottlenecks. The Journal of Heredity, 89, 238–247.

Luikart G, Cornuet J-M (1998) Empirical evaluation of a test for identifying recently bottlenecked populations from allele frequency data. Conservation Biology, 12, 228–237.

Marriage TN, Hudman S, Mort ME et al. (2009) Direct estimation of the mutation rate at dinucleotide microsatellite loci in *Arabidopsis thaliana* (Brassicaceae). Heredity, 103, 310–317.

Muller M-H, Latreille M, Tollon C (2010) The origin and evolution of a recent agricultural weed: population genetic diversity of weedy populations of sunflower (*Helianthus annuus* L.) in Spain and France. Evolutionary Applications, 4, 499–514.

Nagylaki T (1998) The expected number of heterozygous sites in a subdivided population. Genetics, 149, 1599–1604.

NASS-USDA (2015) Adoption of genetically engineered crops in the United States. National Agricultural Statistics Service Report, June Agricultural Survey for years 2000–2015.

Nei M, Maruyama T, Chakraborty R (1975) The bottleneck effect and genetic variability in populations. Evolution, 29, 1–10.

Oerke EC (2005) Crop losses to pests. The Journal of Agricultural Science, 144, 31–14.

Orsini L, Schwenk K, De Meester L et al. (2013) The evolutionary time machine: using dormant propagules to forecast how populations can adapt to changing environments. Trends in Ecology & Evolution, 28, 274–282.

Paetkau D, Calvert W, Stirling I, Strobeck C (1995) Microsatellite analysis of population structure in Canadian polar bears. Molecular Ecology, 4, 347–354.

Paetkau D, Slade R, Burden M, Estoup A (2004) Genetic assignment methods for the direct, real-time estimation of migration rate: a simulation-based exploration of accuracy and power. Molecular Ecology, 13, 55–59.

Parker IM, Rodriguez J, Loik ME (2003) An evolutionary approach to understanding the biology of invasions: local adaptation and general-purpose genotypes in the weed *Verbascum thapsus*. Conservation biology, 17, 59–72.

Peakall R, Smouse PE (2012) GenAlEx 6.5: genetic analysis in Excel. Population genetic software for teaching and research—an update. Bioinformatics, 28, 2537–2539.

Peery MZ, Kirby R, Reid BN et al. (2012) Reliability of genetic bottleneck tests for detecting recent population declines. Molecular Ecology, 21, 3403–3418.

Pimentel D, Zuniga R, Morrison D (2005) Update on the environmental and economic costs associated with alien-invasive species in the United States. Ecological economics: the journal of the International Society for Ecological Economics, 52, 273–288.

Piry S, Alapetite A, Cornuet JM et al. (2004) GENECLASS2: A software for genetic assignment and first-generation migrant detection. The Journal of heredity, 95, 536–539.

Piry S, Luikart G, Cornuet J-M (1999) BOTTLENECK: a program for detecting recent effective population size reductions from allele data frequencies. The Journal of Heredity, 90, 502–503.

Powles SB (2008) Evolved glyphosate-resistant weeds around the world: lessons to be learnt. Pest management science, 64, 360–365.

Rousset F (2008) genepop’007: a complete re-implementation of the genepop software for Windows and Linux. Molecular Ecology Resources, 8, 103–106.

Stewart NC, Warwick SI (2005) Crops Come from Wild Plants — How Domestication, Transgenes, and Linkage Together Shape Ferality. In: Crop Ferality and Volunteerism (ed Gressel J), pp. 9–30. CRC Press, 10.1201/9781420037999.ch2.

Tehranchian P, Riar DS, Norsworthy JK et al. (2015) ALS-resistant mallflower umbrella sedge (*Cyperus difformis*) in Arkansas rice: physiological and molecular basis of resistance. Weed Science, 63, 561–568.

Thomann M, Imbert E, Engstrand RC, Cheptou PO (2015) Contemporary evolution of plant reproductive strategies under global change is revealed by stored seeds. Journal of Evolutionary Biology, 28, 766–778.

Van Etten ML, Chang S-M, Baucom RS (2015) Reduced seed viability as a cost of glyphosate resistance in an agricultural weed. In review, 1–24. http://dx.doi.org/10.1101/030833

Vigueira CC, Olsen KM, Caicedo AL (2013) The red queen in the corn: agricultural weeds as models of rapid adaptive evolution. Heredity, 110, 303–311.

Waselkov KE, Olsen KM (2014) Population genetics and origin of the native North American agricultural weed waterhemp (*Amaranthus tuberculatus;* Amaranthaceae). American Journal of Botany, 101, 1726–1736.

Zar JH (1996) Biostatistical analysis. Prentice Hall, Upper Saddle River.

Carlsson, J. (2008). Effects of microsatellite null alleles on assignment testing. Journal of Heredity. 99(6): 616–623,

Dabrowski, M.J, et al. (2014). Reliability assessment of null allele detection: inconsistencies between and within different methods. Molecular Ecology Resources, 14:361–373.

Foll, M., and Gaggiotti, O (2008). A genome-scan method to identify selected loci appropriate for both dominant and codominant markers: a Bayesian perspective. Genetics, 180(2):977–993.

Foll, M, Fischer, M.C., Heckel, G. Excoffier, L. (2010). Estimating population structure from AFLP amplification intensity. Molecular Ecology, 19:4638–4647.

Peakall, R. and Smouse P.E. (2012) GenAlEx 6.5: genetic analysis in Excel. Population genetic software for teaching and research-an update. Bioinformatics 28, 2537–2539.

Rousset, F. (2008). genepop’007: a complete re-implementation of the genepop software for Windows and Linux. Molecular Ecology Resources, 8:103–106.

Rice, WR (1989). Analyzing tables of statistical tests. Evolution, 43(1):223–225.

Van Oosterhout, C, Hutchinson, WF, Wills, DP, and Shipley, P (2004). MICROCHECKER: software for identifying and correcting genotyping errors in microsatellite data. Molecular Ecology Notes, 4(3):535–538.

